# Integrating predator energetic balance, risk-taking behavior and microhabitat in functional response to untangle indirect interactions in a multispecies vertebrate community

**DOI:** 10.1101/2024.12.20.629406

**Authors:** Andréanne Beardsell, Frédéric Dulude-De-Broin, Gilles Gauthier, Dominique Gravel, Pierre Legagneux, John P. DeLong, Dominique Berteaux, Joël Bêty

**Affiliations:** Département de biologie and Centre d’études nordiques, Université Laval, Québec, Québec, Canada; Département de biologie and Centre d’études nordiques, Université de Sherbrooke, Sherbrooke, Québec, Canada; School of Biological Sciences, University of Nebraska- Lincoln, Lincoln, Nebraska, USA; Chaire de recherche du Canada en biodiversité nordique, Centre d’études nordiques and Centre de la science de la biodiversité du Québec, Université du Québec à Rimouski, Rimouski, Québec, Canada

**Keywords:** Multispecies functional response, predator-prey interactions, predation sequence, energetic, apparent mutualism, prey refuges, Arctic tundra

## Abstract

1. Predator-prey interactions in natural communities are complex, with predators often exploiting multiple prey types and generating indirect interactions among them. Ecological theory has traditionally modeled these interactions using functional responses models which are based on foraging rates, not energy transfers. This approach overlooks how the energy acquisition rate of a predator can alter its behavior and, in turn, the strength of species interactions.
2. Here, we integrate predator energetics into a functional response model to represent trade-offs predators face when foraging on prey that vary in risk and abundance across heterogeneous landscapes. We compared model predictions to 20 years of prey species density and reproductive success data. The mechanistic model was parameterized for an Arctic tundra vertebrate community, where the Arctic fox feeds on cyclic lemmings and eggs of sandpipers (non-risky prey) and gulls (risky prey that often nest in partial refuge like islands). In this system, predator-mediated interactions generate apparent mutualism between lemmings and birds, but its strength varies between species, and the mechanisms underlying this interaction remain unclear.
3. We found that fox energetic balance was highly related to lemming density, with a threshold of 89 lemmings per km^2^ required for a positive energetic balance. Model-predicted gull nest acquisition rates were lowest on islands when the energetic balance of foxes was positive, and highest for nests on the shore when foxes were in deficit. The model that incorporated predator risk-taking behavior and energetic balance produced variation in gull hatching success that most closely matched empirical observations.
4. We documented for the first time that a shift in predator energetic balance, triggering changes in attack and capture probabilities on a risky prey, can be a key mechanism underlying the apparent mutualism between lemmings and gulls. In contrast, for non-risky prey, the indirect effect can be essentially driven by changes in predator movement. These findings highlight how prey characteristics can lead to different mechanisms behind similar indirect interactions.
5. Taken together, our results indicate that mechanistic models integrating species traits, landscape features, and energy-dependent behavioral adjustments can improve our ability to quantify interaction strengths in natural communities.

## 2 Introduction

Natural ecosystems are characterized by a large number of species involved in many predator-prey interactions of variable strength (Berlow et al., 2004; Wootton and Emmerson, 2005). To quantify the strength of biomass and energy flows through ecological communities, classical ecological theory has modeled predators-prey interactions as outcomes of density-driven changes in predator acquisition rates, namely the functional response (Holling, 1959*a*; Murdoch, 1969; Solomon, 1949). While this framework has provided valuable insights, functional responses are based on foraging rates rather than energy transfers. Integrating predator energetics into functional response models can enables us to account the trade-offs predators face when foraging on prey species that vary in risk and abundance across heterogeneous landscapes. Such integration is needed to model multi-prey food webs that characterize the most common ecological communities in nature (Marleau et al., 2020; Portalier et al., 2019; Prokopenko et al., 2023).

Predator–prey interactions are the culmination of a sequence of events that begins with spatial overlap between predator and prey, followed by encounter, detection, attack, and capture (Fig. 1). Probabilities and durations associated with each component allow the formulation of the functional response (Holling, 1959*b*; Lima and Dill, 1990; Wootton et al., 2021). Predator saturation and prey switching are the core concepts of the functional response theory (Holling, 1959*b*; Murdoch, 1969). However, the predation sequence can be modulated by several other factors, including prey characteristics (e.g. presence of defensive structure), the presence and abundance of other prey species (Chan et al., 2017), predator energetic balance, and landscape characteristics such as refuges (Forrester and Steele 2004, Fig. 1). A change in any of these factors can alter the predation sequence and ultimately influencing population dynamics (Beardsell et al., 2023). A better integration of these complexities in mechanistic models of predation in natural systems can greatly increase our understanding of predator-prey interactions and enhance our ability to predict ecosystem responses in a changing world (Cherif et al., 2024; Guiden et al., 2019; Prokopenko et al., 2023; Suraci et al., 2022; Wootton et al., 2021).

**Figure 1.**
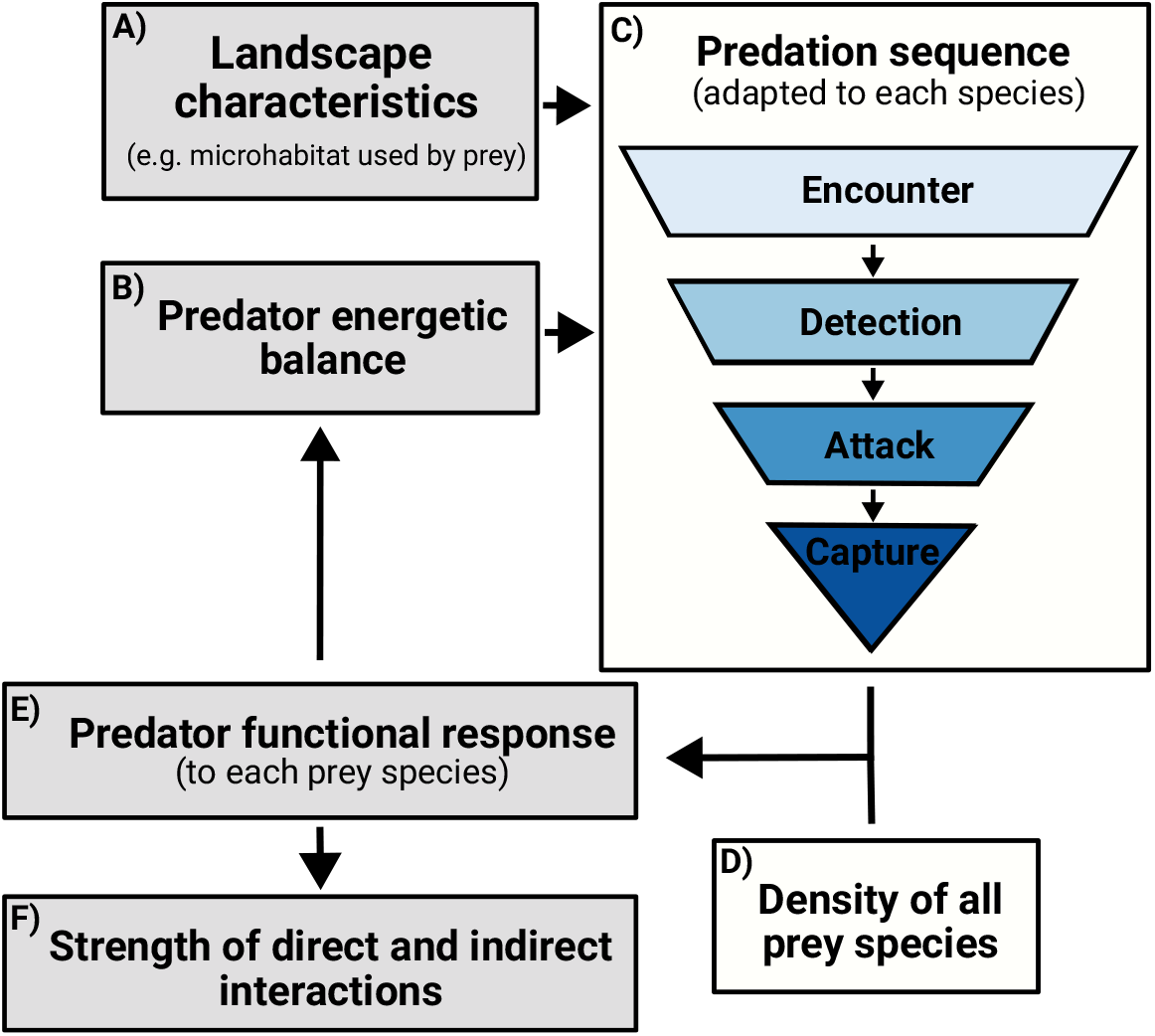
Schematic diagram of the mechanistic model including the effects of landscape characteristics (**A**) and the predator energetic balance (**B**) on the predation sequence (**C**). The quantitative outcome of the predation sequence in relation to the prey density (**D**), namely the functional response (**E**), enables calculation of the energetic balance of the predator, which in turn can influence components of the predation sequence (through changes in predator risk-taking behavior). Finally, a change in the functional response can modulate the strength of direct and indirect interactions (**F)**.

Predators typically forage on diverse prey species, with trade-offs between energy gain and risk of mortality and injury. These can arise from the physical context (steep slopes, strong currents, thorny vegetation) as well as from prey defensive structures (horns, beaks, claws, quills, teeth) combined with aggressive behavior (mobbing, biting, kicking) (see Mukherjee and Heithaus (2013) for review). Predator foraging behavior can vary with the risk of injury, and their energetic state can modulate the propensity to take risks (e.g. Berger-Tal et al. 2009; Embar et al. 2014; Perlman and Tsurim 2008). This short-term behavioral adjustment can be particularly pronounced in endothermic predators, as they generally cannot reduce their metabolic rate to conserve energy when prey abundance is low unlike ectotherms (Kiørboe et al., 2018). Risky foraging activities should be avoided when predators meet their energetic needs (Embar et al., 2014). In contrast, predators facing energetic deficits may take greater risks, as the probability of mortality from not acquiring enough prey increases (Blecha et al., 2018). For instance, European starlings (*Sturnus vulgaris*) increase their attack rates on chemically defended insect larvae when their energetic state (body masses and fat stores) is reduced (Barnett et al., 2007). Such mechanisms could strongly modulate interaction strength, but they have been rarely integrated in predator-prey models (see Prokopenko et al. 2023).

The short-term positive indirect effect of lemmings on tundra nesting birds, mediated by shared predators (apparent mutualism), has been recognized for decades (Summers et al., 1998; Underhill et al., 1993). During the summer, the Arctic fox (*Vulpes lagopus*) is the main predator of lemmings and various bird species (mostly eggs and juveniles) in many ecosystems (Angerbjörn et al., 1999; Giroux et al., 2012). Like many other animals (Wall, 1990), Arctic foxes generally acquire more prey than they immediately consume (Careau et al., 2007). Food hoarding behavior reduces handling time associated with digestion and satiety, resulting in a functional response that is not limited by handling time at high prey densities (Beardsell et al., 2021). Thus, predator saturation has a negligible impact on the indirect interaction between lemmings and nesting birds (Beardsell et al., 2022).

The indirect effect of lemmings on birds varies depending on their traits and the microhabitats they use (Duchesne et al., 2021). For non-risky prey such as passerines and sandpipers, a reduction in Arctic fox daily activity time and distance traveled at high lemming densities can be a main driver of this indirect interaction (Beardsell et al., 2022). In contrast, risky prey such as glaucous gulls (*Larus hyperboreus*) actively defend their nests by harassing potential predators from a distance and diving at them to deter attacks (Clode et al., 2000), posing a potential risk for serious eye, skull, or limb injury to foxes. The risk of injury could be especially pronounced for gulls nesting on islands, which act as partial refuges from foxes for some nesting birds (Corbeil-Robitaille et al., 2024; Duchesne et al., 2021). Foxes typically avoid crossing water to access bird nests (Lecomte et al., 2008), and some types of island micro-habitat could provide a defensive advantage to dangerous prey. Consequently, hatching success of gulls is much higher on small islands than on shores (Gauthier et al., 2015). Thus, the interplay between microhabitats used by nesting gulls and changes in fox risk-taking behavior, triggered by changes in their energetic balance, could generate the strong variation observed in gull breeding success.

We hypothesize that the energetic balance of the predator influences risk-taking behavior and therefore interaction rates between prey species. We formalize our hypothesis with a mechanistic model of the Arctic tundra vertebrate community. Beardsell et al. (2023, 2022) previously developed a mechanistic model of predation in a multiprey system in the Arctic, grounded in detailed empirical observations from a long-term ecological study (Gauthier et al., 2024). Here, we extend this model to integrate predator risk-taking behavior, predator energetic balance, and the presence of prey refuges in the landscape. We assume that microhabitats used by nesting gulls (islands vs shore) and prey defensive behavior modulate attack and capture probabilities of foxes. We also assume that fox risk-taking propensity should be greater when the energetic balance of the predator is negative. We included these two main assumptions into the model and simulated multispecies predator-prey dynamics at varying prey densities. Finally, we confronted model predictions with long-term data on Arctic wildlife densities and demographic parameters. While we lack some empirical data to fully parameterize the model, it extends the framework proposed by Cherif et al. (2024) to a multispecies community in the presence of a risky prey and provides an integrated mechanistic framework to quantify the components of interaction strengths in the wild. Our study provides evidence that the energetic balance of predators can have a significant effect on direct interaction rates and thereby contribute to indirect interactions in complex food webs.

## 3 Methods

### 3.1 Study system

We built the model using detailed empirical data from a long-term ecological study on Bylot Island, Nunavut, Canada (Gauthier et al., 2024). The monitoring area for lemmings, sandpipers, and gulls is located in a 35 km^2^ portion of the Qarlikturvik Valley (73^*°*^ 09’N; 79^*°*^ 56’W). The valley is dominated by a mosaic of mesic tundra, wet polygons, small lakes, and ponds. Wetlands are predominantly used by nesting glaucous gulls, cackling geese (*Branta hutchinsii*), red-throated loons (*Gavia stellata*), and occasionally snow geese (*Anser caerulescens*). Snow geese nesting in the valley are generally associated with nesting snowy owls (*Bubo scandiacus*), whose presence reduces predation by foxes (Bêty et al., 2001). Ground-nesting birds present include passerines (Lapland longspur, *Calcarius lapponicus*), long-tailed jaegers (*Stercorarius longicaudus*) and sandpipers (primarily Baird’s Sandpiper (*Calidris bairdii*) and White-rumped Sandpiper (*Calidris fuscicollis*)). Two species of small mammals are present: brown (*Lemmus trimucronatus*) and collared (*Dicrostonyx groenlandicus*) lemmings (Gauthier et al., 2013). The brown lemming is the most abundant species and shows large-amplitude fluctuations of abundance every 3-5 years, while collared lemmings show weak-amplitude population fluctuations (Fauteux et al., 2015; Gruyer et al., 2008). The nest density of gulls and sandpipers in the study area was relatively constant between years (Beardsell et al., 2022; Gauthier et al., 2015). Some species nesting at low densities (e.g., loons and cackling geese) were not included in the model, as their presence is unlikely to significantly impact the energetic balance of foxes. The Arctic fox is an active searching predator that travels long distances daily within its territory during the summer (Poulin et al., 2021).

### 3.2 Multiprey mechanistic model of predation

The functional response model, inspired by the theoretical framework of Holling (1959*b*) and Wootton et al. (2021), was derived by breaking down predation events into 4 steps: (1) encounter, (2) detection, (3) attack and pursuit, (4) capture and manipulation (Fig. 1C). At the core of this modular framework is the adaptation of the model to different prey species according to their antipredator behavior and the predator hunting behavior (Beardsell et al., 2021, 2022; Cherif et al., 2024; Wootton et al., 2021). We built upon the mechanistic model of predation by Arctic foxes on lemmings and sandpiper nests previously developed in the same study system (Beardsell et al., 2021, 2022). Remarkably, functional responses derived from the model were consistent with field observations for various prey species, including lemmings and the nests of geese, passerines, and sandpipers (Beardsell et al., 2021). Here, we expand the model to include predation on glaucous gull nests. Figure 2A provides an overview of the multiprey model including the three prey species and of the mechanisms involved.

**Figure 2.**
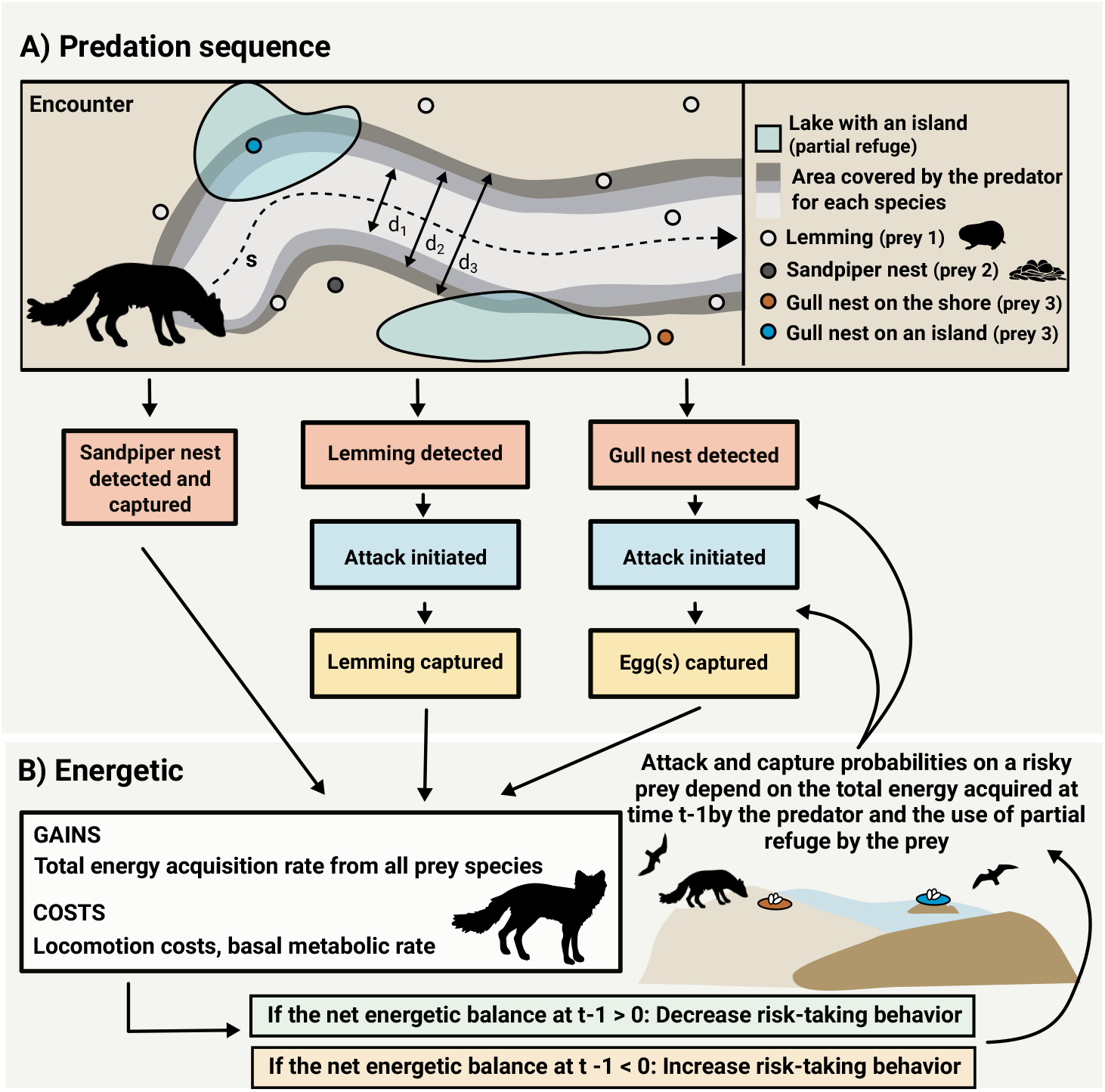
**A)** Conceptual mechanistic model of Arctic fox functional response to density of lemmings, sandpiper nests (non-risky prey), and glaucous gull nests (risky prey). Each box represents one or more components of predation (search, prey detection, attack decision, and capture). Arrows represent the probability that the predator reaches the next component. **B)** Schematization of the integration of the energetic component in the predation model. The attack and capture probabilities of a gull nest by the Arctic fox vary according to whether the nest is located in a partial refuge and the net energetic balance of the predator at time t-1.

An attack by the predator on a gull nest (prey 3) can occur when the predator is at a distance (*d*_3_), defined as the maximum distance at which the predator will attack a nest (in 2D, detection region = 2*d*_3_; Pawar et al. 2012). As gulls are highly conspicuous, we assumed that within the attack distance, the detection probability is 1. As not all nests within the searched area may be attacked and captured by the predator, we introduced the attack and capture probabilities. Attack and capture probabilities at time t are determined by the daily net energetic balance of the predator at time t-1 (*E*_*net*_(*N*_1_, *N*_2_, *N*_3_)_*t−*1_, Fig. 3). The next section describes the calculation of *E*_*net*_(*N*_1_, *N*_2_, *N*_3_)_*t−*1_. Capture efficiency of gull nests located on islands at time *t* (*α*_3,*is*_; km^2^ day^*−*1^) is thus expressed as the product of the distance traveled by the predator when active (s; km day^*−*1^, which depends on lemming density *N*_1_ (see below)), the attack distance (*d*_3_; km), and attack and capture probabilities (*f*_3,3,*is*_, function of *E*_*net*_(*N*_1_, *N*_2_, *N*_3_)_*t−*1_):

**Figure 3.**
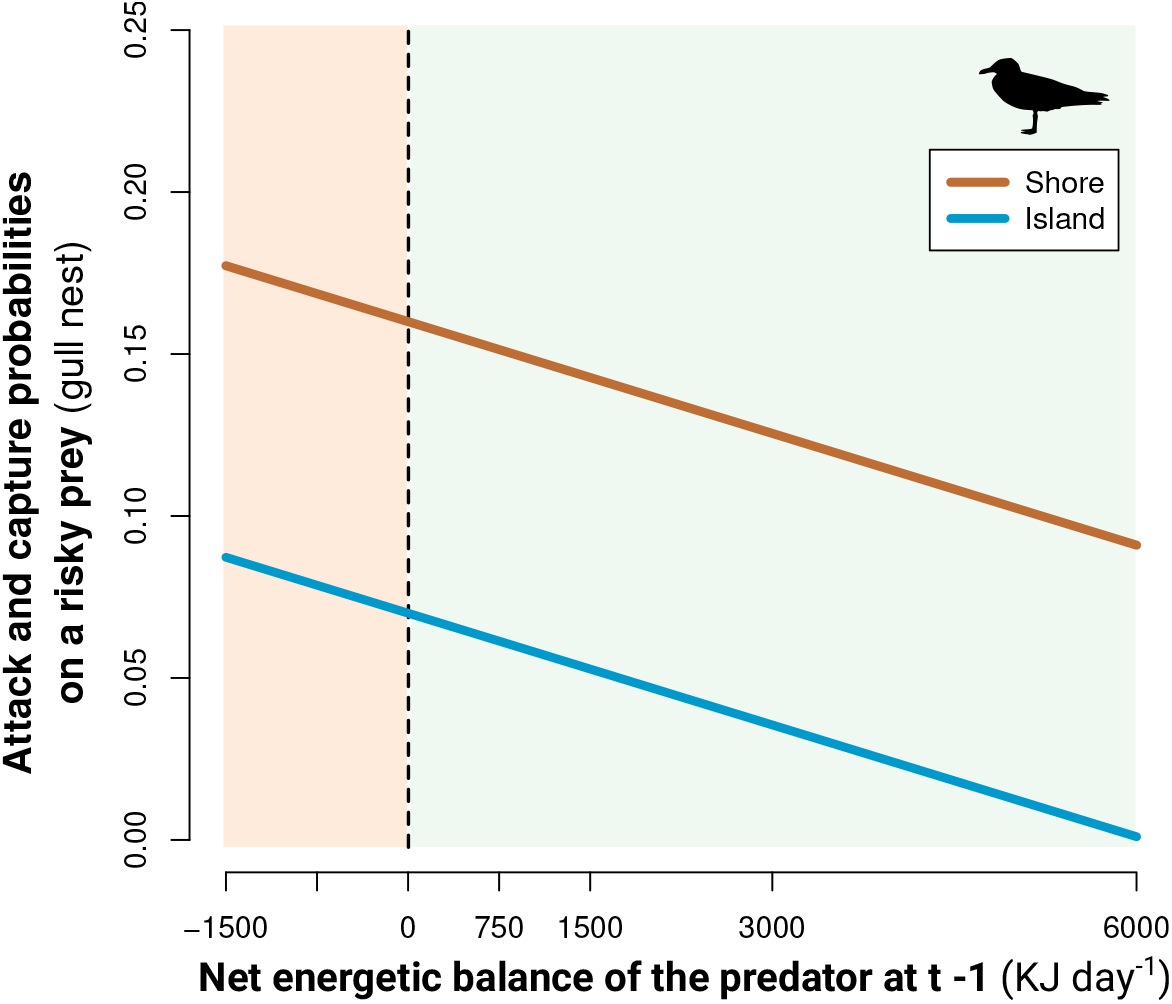
Hypothetical relationships between daily attack and capture probabilities on a risky prey (gull nest) by the Arctic fox at time t as a function of the net energetic balance of the predator at time t-1. Attack and capture probabilities of a nest located on the shore (orange line) are expected to be higher than those of a nest located in a partial refuge (island; blue line). Areas where the net energetic balance is >0 are in green and <0 in orange.

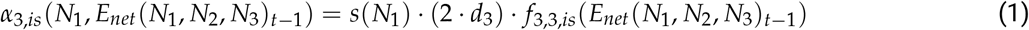

An equivalent equation for nests on the shore can be obtained by substituting all indices *is* (nests on islands) with *sh* (nests on the shore), making the attack and capture probabilities microhabitat dependent.

The number of gull nests acquired (nests per fox day ^*−*1^) is expressed as the sum of gull nests acquired from both habitats:

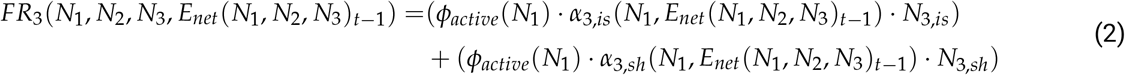

Where *ϕ*_*active*_ is the proportion of time the predator is active in a day and *N* is the density of each prey (prey item km^*−*2^). The values of *ϕ*_*active*_ and *s* are correlated and are a function of lemming density (*N*_1_). This function was partially obtained from empirical data (GPS-tracking of 16 foxes (7 females and 9 males) during the summers 2018 and 2019; Fig. S9, Beardsell et al. 2022). Changes in Arctic fox movement behaviors (*ϕ*_*active*_ and *s*) have been identified as important mechanisms driving the positive effects of lemmings on non-risky prey, such as sandpipers, generating up to 31% variation in their absolute nesting success (Beardsell et al., 2022). Gull nest density (*N*_3_, nests km^*−*2^) is calculated for each microhabitat (*N*_3,*sh*_, *N*_3,*is*_). At the study site, 75% of nests were located on islands (*n* = 264 nests, from 2004 to 2023). Therefore, we introduced the probability that a nest is located on an island (*p*_*r*_) to distribute the overall nest density, *N*_3_, measured in the study area between island and shore nests:

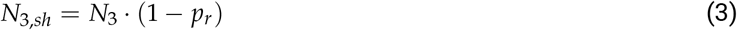

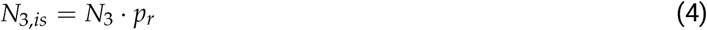

For simplicity, prey handling was not included in the predation model, as these processes (time associated with pursuit and manipulation per prey item) play a minor role in our system (Beardsell et al., 2021, 2022). Very similar results were obtained with predation models including prey handling parameters. As Arctic foxes can capture more prey than they consume in the short-term (Careau et al., 2008; Samelius and Alisauskas, 2000), prey digestion time was not included in the model as a potential limit on acquisition rate. For more details on construction of the model for sandpipers and lemmings, see Beardsell et al. (2022). The next section describes the integration of the predator energetic balance into the model.

### 3.3 Integrating energetics into the predation model

The daily energy gain rate for an individual fox consuming any prey type (*ER*_*i*_(*N*_1_, *N*_2_, *N*_3_, *E*_*net*_(*N*_1_, *N*_2_, *N*_3_)_*t−*1_)) is expressed as the product of the predator acquisition rate (for instance Eq. 2 for prey 3) and the energetic content of a prey item (*E*_*i*_, KJ per prey item). The net energy gain rate of all prey acquired (*E*_*net*_(*N*_1_, *N*_2_, *N*_3_); per fox day^*−*1^) is computed as:

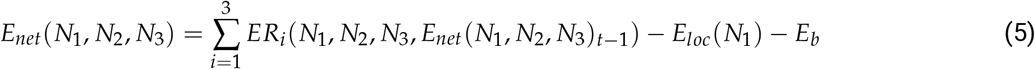

where the total energy gain rate of all prey species is reduced by the energetic costs of locomotion (*E*_*loc*_; KJ day^*−*1^) and basal metabolic rate (*E*_*b*_; KJ day^*−*1^). Considering that movement is a substantial portion of daily energy expenditure in canids (Garland, 1983; Gorman et al., 1998) and that handling time of all three prey species is very low (e.g., 250 sec. to consume or hoard a sandpiper nest and 125 sec to pursue, consume, or hoard a lemming; Beardsell et al. 2022), we assumed that short-term energy costs associated with the time spent pursuing prey (including running and swimming) and handling prey are negligible, and that a portion of these costs is already included in locomotion costs.

The net energetic costs of locomotion (*E*_*loc*_; KJ day ^*−*1^) was estimated by the product of the minimum net costs of transport per kilogram per unit distance (*MCOT*_*min*_, KJ kg^*−*1^ km^*−*1^), the average predator body mass (*M*; kg), the predator speed (km day^*−*1^; *s*) and proportion of time spent active (*ϕ*_*active*_):

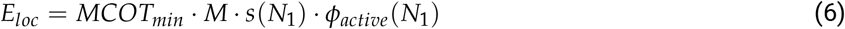

*MCOT*_*min*_ was estimated using the following allometric equation from Taylor et al. (1982):

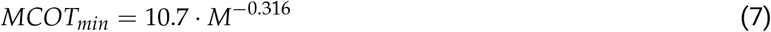

This equation was derived using an energy conversion of 20.1 J ml^1^ *O*_2_ and assumes a negligible contribution from anaerobic glycolysis (Taylor et al., 1982). Additionally, we include only the slope of the equation from Taylor et al. (1982) to compute the incremental cost of locomotion because basal metabolic rate is accounted for separately (Halsey, 2016). We estimated the net energetic cost of locomotion at 23.8 KJ km^*−*1^ for an average fox body mass (3.2 kg), which is consistent with an estimate from the kit fox (*Vulpes macrotis*) in natural settings (15.6 kJ km^*−*1^; Girard 2001). The net energetic cost of locomotion estimated here is also consistent with Fuglei and Oritsland (2003), who used indirect calorimetry to measure the metabolic rate of an Arctic fox running on a treadmill. From their reported values in July, we calculated an energetic cost of 27.4 KJ km^*−*1^ for an average body mass of 3.2 kg.

To estimate the basal metabolic rate (*E*_*b*_), we used the equation presented by Careau et al. (2008) based on Arctic fox body mass:

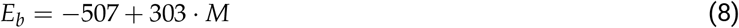

Which leads to an average basal metabolic rate of 472 KJ per day. Parameter values of the energetic model are summarized in Table 2, and details on the estimation of parameter values can be found in Supplementary Methods.

### 3.4 Parameter values

Behavioral processes affecting the interaction between foxes and gulls are relatively well documented, although the parameters needed to fully define the relationship between predator energetic balance and its attack and capture probabilities of gull nests are largely unknown. We inferred parameter values based on the following considerations. First, unlike passerines and sandpipers, gulls actively defend their nests with dive attacks, posing a potential risk of serious eye, skull, or limb injury to foxes (see Fig. S8 for images of interactions between gulls and foxes). Therefore, we assume that attack and capture probabilities of foxes at time t decrease linearly with the daily net energy acquired by the predator at time t-1 (Fig. 3). Specifically, attack and capture probabilities should be higher when the predator energetic balance is negative (increased risk-taking behavior) and lower when the energetic balance is positive (reduced risk-taking behavior, Fig. 2B).

Second, because foxes attacking island nests must swim, they cannot achieve the same speed or adopt the same attack and defensive positions as they would on the shore when harassed by gulls. Therefore, we assume that both attack and capture probabilities are lower for nests on islands (Fig. 3). An experimental study at our site shows that daily fox attack and capture probabilities, measured with artificial nests (537 nests), are 4-11 % lower for nests on islands than on the shore (Beaudoin, 2025).This supports our assumption and justifies the difference in intercept values used in Fig. 3. For Greater Snow Geese, a larger species than gulls (body mass ranging from 1.6 to 3.3 kg vs. 1.25 to 2.7 kg for gulls) that also defends nests, we estimated fox attack and capture probabilities at 0.005 based on 124 hours of direct behavioral observations (Beardsell et al., 2021). This value is the lowest attack and capture probabilities used in the model for gulls (nests on islands when the fox is at its highest positive energetic balance). Given the limited empirical data to support this relationship, we generated six different functions, both linear and sigmoidal (Fig. S7). The model outputs obtained with one function are presented in the results, but results from other simulations are presented in Fig. S7B1 and B2.

For island nests, characteristics such as water depth and distance to the shore could influence fox attack and capture probabilities. However, in our system, gulls select islands that are farther from shore and surrounded by deeper water (Corbeil-Robitaille et al., 2024). Among the islands used, Beaudoin (2025) found that distance to shore had only a negligible effect on nest survival, while water depth had no effect. These findings justify the use of fixed parameter values for island nests in our model. All parameter values used in the predator model for gulls and other prey (lemmings, prey 1, and sandpiper, prey 2) and their source can be found in Supplementary methods.

### 3.5 Estimating gull hatching success from the mechanistic model

The number of prey items acquired by the predator population per day for each prey species (*i* = 1,2,3) is given by the product of the functional response and the predator density (*NR*, ind. km^*−*2^):

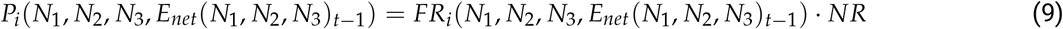

In the study area, the home ranges of Arctic foxes are generally stable throughout the summer. Thus, predator density was estimated based on average home range size (18.2 km^2^ based on 57 home ranges from 2008 to 2016) and the average proportion of overlap (0.18; see Beardsell et al. 2023; Dulude et al. 2023 for more details). Eq. 9 allowed us to estimate the annual hatching success of gulls (prey 3) for the entire range of lemming density measured at the study site. For each value of lemming density, we calculated the number of nests acquired for 28 days (the average duration between the gull laying date and hatching date), while considering that the density of nests decreases each day due to predation by foxes. To estimate hatching success, we assumed that the nesting period of gulls occurs simultaneously across individuals, that fox predation is the only cause of nest failure, and that predated nests are not replaced.

### 3.6 Model evaluation with field observations

We evaluated the model output (gull hatching success) using 20 years (2004 to 2023) of glaucous gull nest monitoring in the study area (Gauthier et al., 2024, 2015). Nests are scattered throughout the landscape and typically located on small islands or on the shore of ponds and small lakes in wetlands. The incubation period of gulls extends from early June to mid-July. We systematically searched the entire study area for gull nests in June of all years. Nests are conspicuous, and gulls typically reveal their presence from a relatively long distance with alarm calls and behavioral displays. Most nests were revisited at 1 to 2 week intervals to check their contents and to look for signs of predation. A nest was considered to have hatched successfully if at least one egg had hatched. We estimated lemming density using live trapping from 2004 to 2023 (see Fauteux et al. 2018 for methods). To obtain an estimate of the total lemming density during the gull incubation period, we summed the density of the two lemming species and averaged the estimates for June and July.

The relationships between hatching success, nest microhabitat (shore, island) and lemming density were fitted using a generalized linear mixed model based on the logistic-exposure method (Shaffer, 2004). The method estimates daily nest survival and explicitly accounts for variation in the length of the nest monitoring period. Exposure time was computed by determining the number of days between the date of nest discovery and one of the following dates: (1) for successful nests, the observed or estimated hatching date or the date of the last visit if no information on hatching date was available, (2) for failed nests, the observed predation date or the midpoint between the date on which fate was determined and the last date the nest contained eggs. Nests with undetermined fate were not included in the analysis. We built a set of candidate models to investigate the effect of lemming density, and nest microhabitat (as fixed effects) on gull hatching sucess (a binary response variable). We included the year as a random variable. Models were ranked according to Akaike’s Information Criterion corrected for small sample size (see Table S3). Since the global model had full support (i.e. an Akaike weight of 1), parameter estimates and 95% confidence intervals were computed from this model. Daily nest survival estimates provided by the statistical model were converted into hatching success (i.e. nest survival over a 28-day nesting period). All analyzes and modeling were conducted in R version 4.4.0 (Team, 2024).

## 4 Results

The mechanistic model indicated that lemmings are the most important contributor to the energy budget of foxes. More importantly, the model showed that the energetic balance of foxes changes from negative to positive at a threshold of 89 lemmings per km^2^ (Fig. 4A). Lemmings constituted 83 and 99% of the daily energy acquired by foxes in a low (50 ind. per km^2^) and a high (650 ind. per km^2^) lemming year, respectively (Fig. 4B). They represented less than 50% of the energy acquired by foxes below approximately 10 lemmings per km^2^. Variation in the density of sandpiper and gull nests (within the variation observed in our system; see Table 1) generated little change in the energetic gain rate of foxes compared to lemmings (346 KJ for gull nests and 417 KJ for sandpiper nests compared to 6652 KJ for lemmings).

**Table 1:** Symbol definition and parameter values used in the multiprey mechanistic model of fox predation as a function of the density of lemmings (prey 1), sandpiper nests (prey 2) and glaucous gull nests (prey 3). Parameter values for foxes, lemmings, and sandpipers, as well as methods used to obtain the values, can be found in Beardsell et al. (2022) unless mentioned otherwise.

| Parameter name | Symbol | Value(s) and unit | Additional information |
| --- | --- | --- | --- |
| **Arctic Fox** |  |  |  |
| Daily proportion of time the predator spent active | $\phi_{active}$ | See Fig. 9 | The value is a function of $N_1$. Based on accelerometry data (23 summer foxes, 2018-2019). |
| Daily distance traveled when active | $s$ | See Fig. 9; km day<sup>-1</sup> | The value is a function of $N_1$ and correlated to $\phi_{active}$. Based on high-frequency GPS data (23 summer foxes, 2018-2019). |
| Average fox density | $NR$ | 0.13 ind. km<sup>-2</sup> | Based on the average fox home range size in the study area (Beardsell et al., 2023). |
| **Lemmings** |  |  |  |
| Lemming density | $N_1$ | 216 ind. km<sup>-2</sup> | Average 216 ind. km<sup>-2</sup> (range from 2 to 710 ind. km<sup>-2</sup>) |
| Maximum reaction distance | $d_1$ | 0.0075 km | Estimated distance from 29 observed attacks (from 1996-1999, 2019) |
| Average detection and attack probability within the reaction distance | $f_{2,1} \cdot f_{3,1}$ | 0.15 | - |
| Capture probability | $f_{4,1}$ | 0.51 | Direct observations of the outcome of 268 attacks (from 1996-2019) |
| **Sandpiper nests** |  |  |  |
| Sandpiper nest density | $N_2$ | 2 nests km<sup>-2</sup> | Average 2 nests km<sup>-2</sup> (range from 0.25 to 4.25 nests km<sup>-2</sup>) from 2005 to 2019. |
| Maximum reaction distance | $d_2$ | 0.085 km | Based on flushing distance (Smith and Edwards, 2018) |
| Average detection probability within the reaction distance | $f_{2,2}$ | 0.029 | Based on flushing probability (Beardsell et al., 2021) |
| **Glaucous gull nests** |  |  |  |
| Gull nest density | $N_3$ | 0.26 nests km<sup>-2</sup> | Average 0.26 nests km<sup>-2</sup> (range from 0.05 to 0.54 nests km<sup>-2</sup>) from 2008 to 2016. |
| Proportion of nest in partial refuge (island) | $p_r$ | 0.75 | From 262 nests monitored from 2004 to 2023. |
| Average clutch size | $CS_3$ | 2.6 | From 175 nests monitored from 2004 to 2023. |
| Reaction distance (attack distance) | $d_3$ | 0.03 km | - |
| Attack and capture probabilities of a nest on the shore | $f_{3,3_{sh}}$ | See Fig. 3A | - |
| Attack and capture probabilities of a nest on an island | $f_{3,3_{is}}$ | See Fig. 3A | - |

**Table 2:**
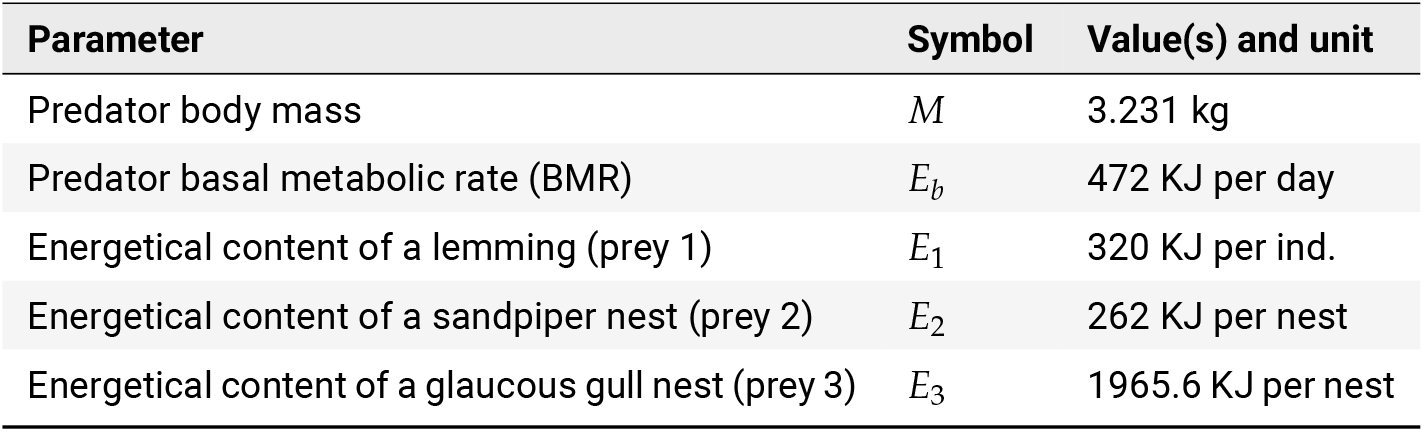
Energetic parameters of the Arctic fox and energetical content of all prey species.

**Table 3:** Variables, number of parameters (K), difference in AICc value between the current model and the preferred model (*δ*AIC_*c*_), Akaike weight (w_*i*_), and log-likelihood (LL) of the candidate models explaining glaucous gull hatching success (*n* = 155 nests) on Bylot Island (Nunavut, Canada) from 2004–2023. We did not included a model with nest microhabitat only due to convergence issues. Year was included as a random effect in all models.

| Variables | K | $\delta AIC_c$ | $w_i$ | LL |
| --- | --- | --- | --- | --- |
| Lemming density, Nest microhabitat | 4 | 0.00 | 1.00 | -106.36 |
| Lemming density | 3 | 68.60 | 0.00 | -141.72 |
| Null | 2 | 73.81 | 0.00 | -145.36 |

**Figure 4.**
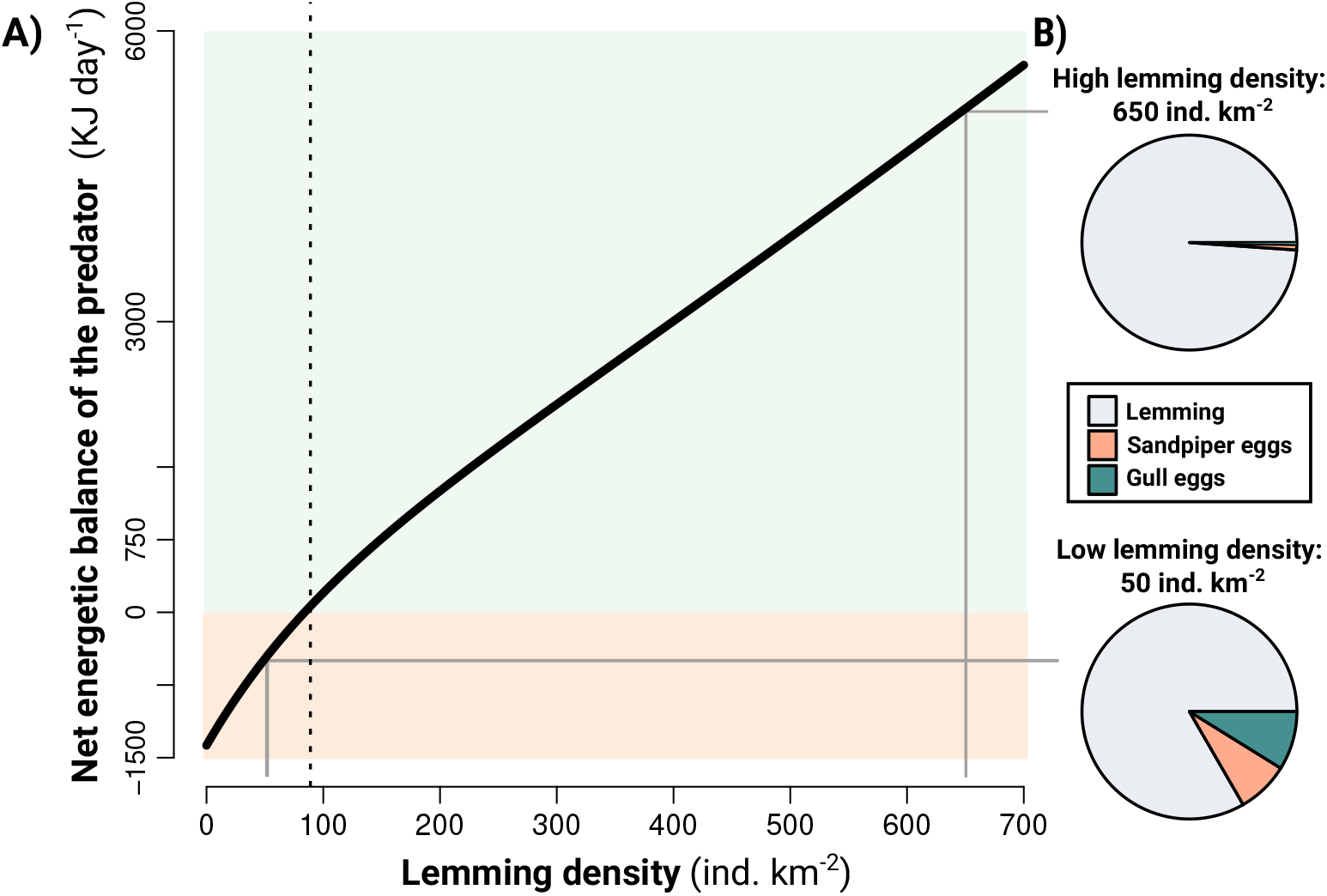
**A)** The black line shows the predator net energetic balance as a function of lemming density for an average-mass fox. Areas where the net energetic balance is >0 are green and <0 are red. The dashed line shows the lemming density threshold at which the net energetic balance of foxes changes from negative to positive. **B)** Pie charts show the relative contribution of the three prey species to the net energetic balance of foxes at high (650 ind. km^*−*2^) and low (50 ind. km^*−*2^) lemming densities. Densities of sandpiper and gull nests are set at average densities (respectively 2 and 0.26 nests km^*−*2^) over the range of lemming density.

The functional responses of the Arctic fox to the risky prey (gull nest) were generated from the multiprey mechanistic model (Fig. 5). The slope of the functional response (the product of capture efficiency (*α*; km^2^ day^*−*1^) and the proportion of time the predator spent active), varied from 0.125 to 0.42 km^2^ day^*−*1^ for nests on the shore (Fig. 5B) and from 0.02 to 0.19 km^2^ day^*−*1^ for nests on islands (Fig. 5A). Acquisition rates of gull nests were lowest for nests on islands when the energetic balance of foxes was positive, and highest for nests on the shore when foxes were in deficit. The variation in predator acquisition rates, generated by the model linking predator risk-taking behavior and energetic balance, is 70% for island nests and 47% for shore nests (Fig. 5). The remaining variation, 30% for island nests and 53% for shore nests, is due to changes in predator movement parameters induced by change in lemming density (Fig. 5). The functional responses of the Arctic fox to a non-risky prey (sandpiper nests) were generated from the model (Fig. S10), with all variation in predator acquisition rates driven by changes in predator movement parameters induced by lemming density.

**Figure 5.**
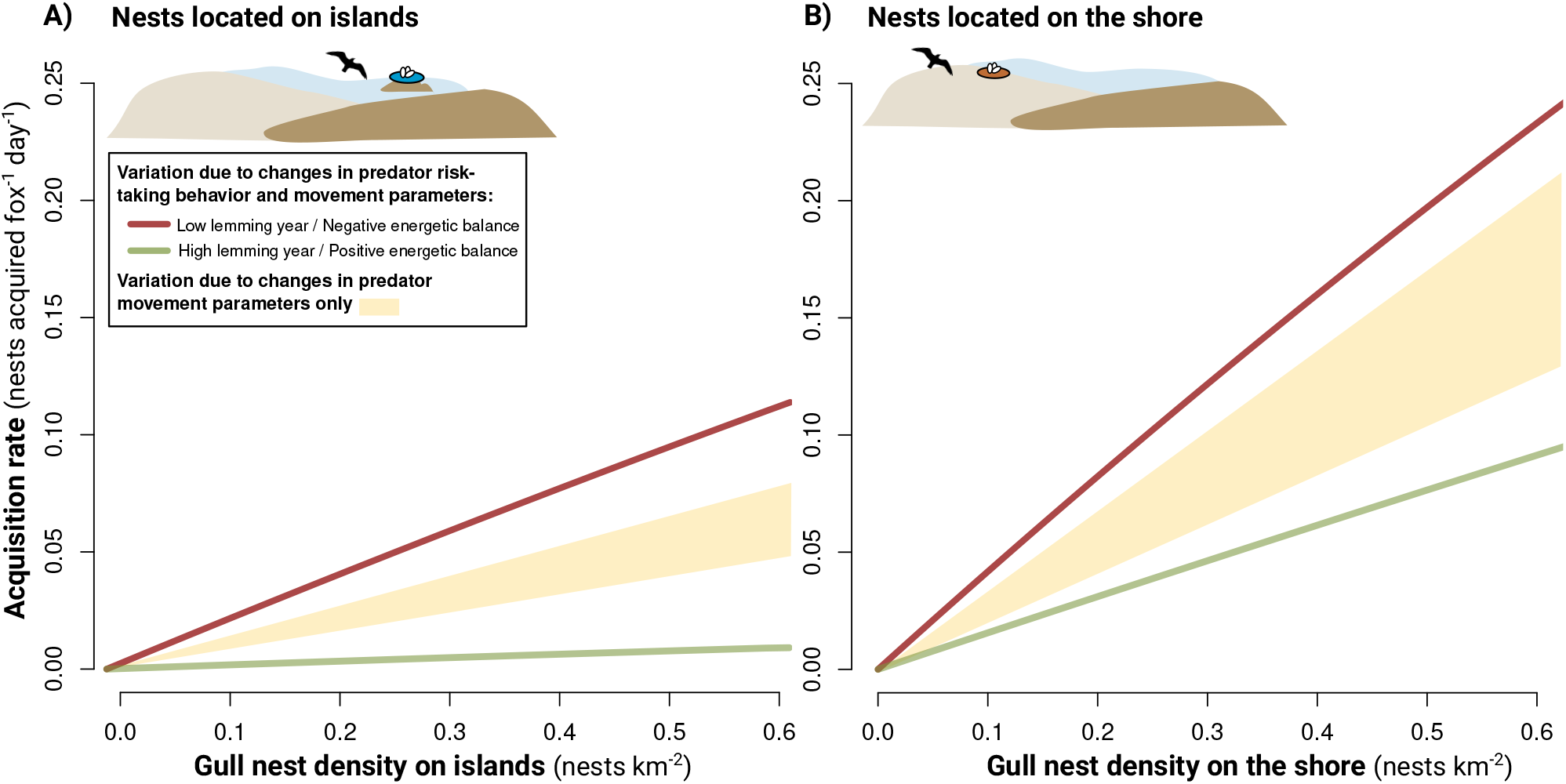
Functional responses of the arctic fox to the risky prey (gull nest) generated from the mechanistic multiprey model for both habitats (islands **(A)** and the shore **(B)**), and for a negative (red lines) and positive (green lines) energetic balance of the predator. Positive and negative energetic balance correspond, respectively, to lemming densities of 650 and 50 ind. per km^2^. The shaded area is based on a model without predator risk-taking behavior, retaining only changes in predator movement parameters (as a function of lemming density; Fig. S9). Sandpiper density was set at the average value of 2 nests per km^2^.

Annual lemming density during the gull incubation period varied from 2 to 710 individuals per km^2^ from 2004 to 2023. A total of 155 glaucous gull nests were monitored until hatch during this period (111 located on islands and 44 on shores). Hatching success was positively related to summer lemming density (*β* = 0.63, 95%CI = 0.25, 1.15) and was lower for nests located on shores compared to islands (*β* = -3.29, 95%CI = -4.29, -2.43, Fig. 6). Variation in gull hatching success generated by the mechanistic model, that incorporated predator risk-taking behavior and energetic balance, aligned with empirical observations, as indicated by the mechanistic model prediction falling within the 95% CI of the empirical statistical model over much of the range in lemming density in both habitats (Fig. 6). For instance, at lemming densities of 50 and 650 ind. per km^2^ (which correspond, respectively, to a negative and positive energetic balance of the predator), the mechanistic model estimated gull hatching probability to vary from 0 to 0.46 on shores and 0.51 to 0.94 on islands, while the statistical model estimated it to vary from 0 to 0.75 on shores and 0.75 to 0.99 on islands. A model without predator-risk taking behavior generated less variation in gull hatching success, especially for island nests, where it generates only 11% of variation (0.64 to 0.75; Fig. 6).

**Figure 6.**
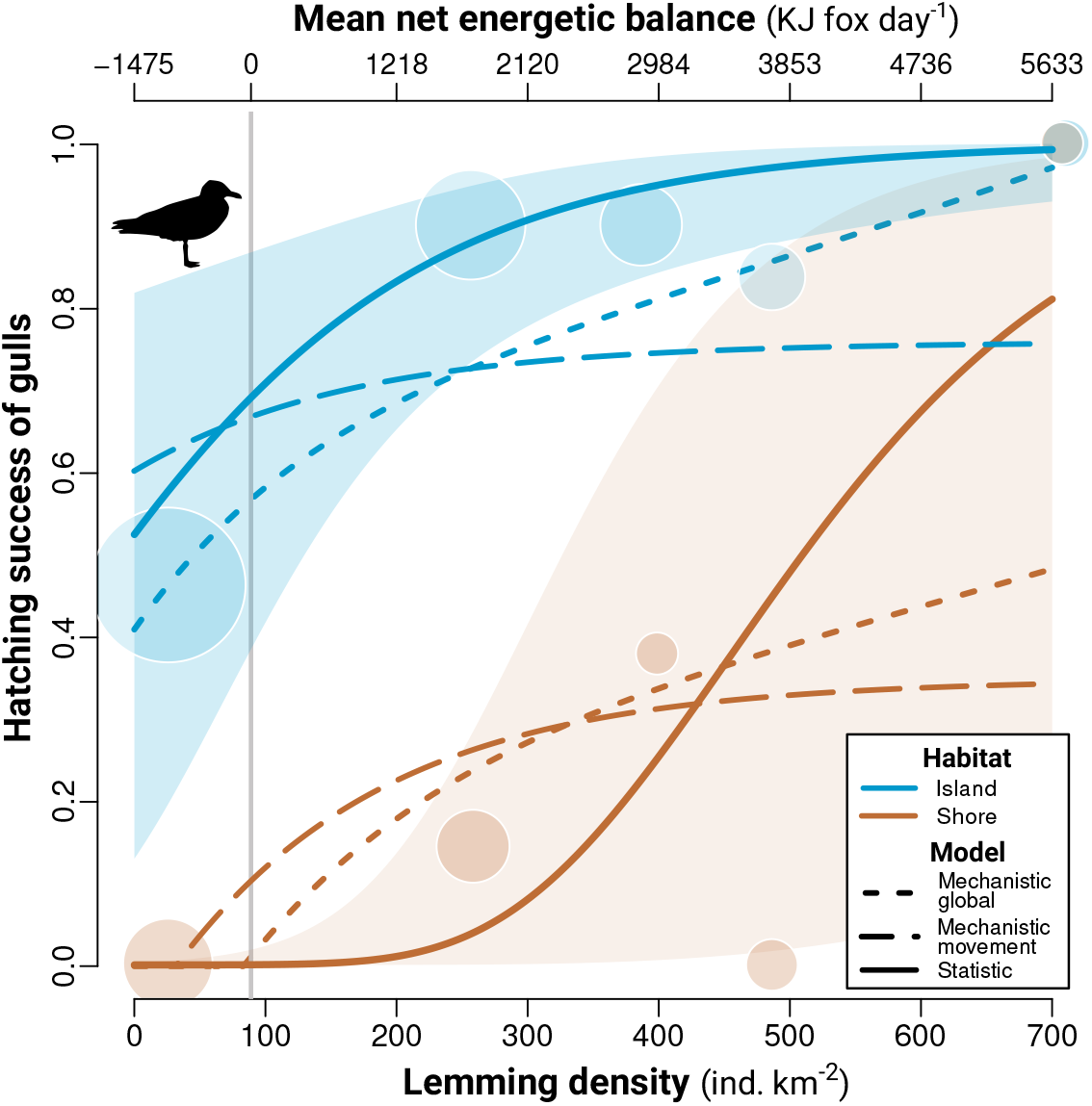
Hatching success probability of glaucous gulls nesting on Bylot Island (Nunavut, Canada) in relation to lemming density and nest microhabitat (shore vs. islands) from 2004 to 2023. The full line is the statistical model prediction over the average duration between the gull laying date and hatching date (28 days) and is presented with its 95% CI. Circles represent empirical hatching success (calculated for a 28-day exposure) averaged over lemming density intervals of equal size and are provided to illustrate model fit with the data. Circle size is proportional to the total number of observations (*n* = 111 nests on islands and 44 on shore). Colored dashed lines show gull hatching success in relation to lemming density and nest microhabitat, based on two mechanistic models. The longest dashed line represents a model without predator risk-taking behavior, retaining only changes in predator movement parameters, while the shortest dashed line shows the global model. The vertical grey line shows the lemming density threshold (predicted by the model) where fox energetic balance changes from negative to positive.

## 5 Discussion

Our study is a first critical step towards the integration of energetic and landscape characteristics into predator-multiprey models. It also presents one of the rare mechanistic models parameterized in a natural community. Such a model can greatly improve our ability to understand the drivers of direct and indirect interaction strengths in complex ecological communities that produce phenomena such as short-term apparent mutualism. By explicitly incorporating the sequence of events associated with the consumption of different prey species by a predator within an energetic framework, our model provides a way to disentangle the interplay between predator energetic balance, risk-taking behavior, and microhabitat heterogeneity. Our results highlight the various mechanisms that can drive indirect interactions among prey and generate strong inter and intra annual variation in prey survival.

Although many empirical studies have linked predator energetic state with risk-taking behavior (Barnett et al., 2007; Blecha et al., 2018; Moran et al., 2021), such relationships are rarely explicitly included in functional response models (Prokopenko et al., 2023). Integration of energy is crucial for understanding interaction strengths, since many species traits and behavior, including movement rate, territoriality, home range size, hunting tactics, and risk-taking propensity, are highly plastic and can be strongly influenced by the energetic balance (e.g., Blecha et al. 2018; Lyon et al. 2018). Our modeling approach is innovative as it integrates (1) habitat-specific multispecies functional responses to estimate predator energy acquisition, and (2) the feedback loops on the predation sequence caused by changes in the predator energetic balance. The approach is transferable to other systems and should benefit from emerging biologging technologies, such as high-frequency GPS tracking, acoustic recording, and accelerometry (Studd et al., 2021; Williams et al., 2014). This modeling approach should strongly enhance our understanding and quantification of interaction strengths in free-ranging animals.

The lemming density threshold for a positive energetic balance of foxes (that is 89 ind. per km^2^) is consistent with previous research conducted at our study site indicating a very low occurrence of fox reproductive activity below 100-170 lemmings per km^2^ (Bergeron et al., 2025; Juhasz et al., 2020). In our model, fox movement parameters (distance traveled and time spent active) were functions of lemming density (see Beardsell et al. 2022). Although these relationships were not the focus of the study, these two parameters, in addition to the risk-taking behavior, are likely to be related to predator energetic balance. Linking energetic balance to key components of predation, such as predator movement, detection, attack and capture probabilities of the prey, could strongly improve our ability to develop general mechanistic models of predation.

Our model indicated that lemmings are the main contributor to the energy budget of Arctic foxes, comprising an average of 92% of their energy gain rate across the observed range of lemming densities. This result is consistent with Angerbjörn et al. (1999) and Elmhagen et al. (2000), who reported that small rodents accounted for an average of 87% of the Arctic fox diet in Siberia and 85% in Sweden based on the scat analysis. In our system, lemming density is highly correlated with predator energetic balance, making it difficult to disentangle their relative importance. However, explicitly modeling energetic balance provides a key advantage for extrapolation to systems where the relationship between prey density and predator energy intake is less direct or predictable. Arctic foxes can persist in environments where small rodents are absent or rare, relying on colonial-nesting birds or carrion (Angerbjörn et al., 1994; Frafjord, 1993). In such systems, the energetic contribution of each prey species may differ substantially. Moreover, prey density does not necessarily translate into energy acquisition—especially for prey that are difficult to capture (e.g., colonial birds with strong nest defense), or occur at low density but offer high energetic gains (e.g., carcasses). By modeling energy acquisition directly, our framework can be applied to other predator–prey systems with varying community compositions, capturing how prey density and profitability shape predator behavior and prey survival—insights missed by models based solely on prey density.

Predators foraging on diverse prey species are common in natural ecosystems. A classic idea in ecology is that predators can stabilize prey populations by adjusting their ‘preferences’ according to the relative abundance of prey (the concept of prey switching; Murdoch 1969). The term ‘preference’ encompasses a wide range of mechanisms, including the probability of detecting, attacking, and capturing a prey item (Chesson, 1984). However, in most cases, the parameter that induces prey switching in predator-prey models does not have an explicit mechanistic interpretation (DeLong, 2021). This parameter is typically integrated through the type III functional response, where the space clearance rate (or capture efficiency) is an increasing function of prey density. One mechanism often suggested for prey switching is predator learning or experience (Real, 1977). Although these cognitive processes can be important, our model illustrates that changes in attack probability on a given prey species also can be driven by changes in the energetic state of the predator per se. Our approach, based on dynamic multispecies functional responses, accounting for total prey densities, allows the integration of energy-dependent mechanisms in predator-multiprey models.

In our model, predator decisions are modulated by short-term energy acquisition rates the previous day. However, energetic balance over a longer time also may play a role in decision-making, particularly if the predator can store energy, either through exogenous or endogenous reserves. This could become especially important when a pulse of abundant prey is available (e.g., a bird colony or carcasses) and used by the predator to cover its energetic needs during periods of low prey availability. Moreover, predators experiencing long-term energy deficits may adjust their spatial behavior, shifting from residency to migration or nomadism. For instance, Arctic foxes can increase extraterritorial excursions during periods of low prey abundance (Lai et al., 2017). Identifying the appropriate time scale to link predation components (such as attack probability) to predator energetic balance is essential, as it could influence predator behavior and interaction strengths.

The positive short-term indirect effect of lemmings on tundra-nesting birds due to shared predators was reported decades ago (Summers et al., 1998; Underhill et al., 1993) and has been studied across the Arctic since then. Changes in the daily activity budget and distance traveled by arctic foxes have been identified as important mechanisms leading to positive indirect effects of lemmings on non-risky prey like sandpipers and passerines (Beardsell et al., 2022). Unlike passerines and sandpipers, gulls can defend their clutches against arctic foxes, and we show that changes in predator energetic balance, translating into a change in attack and capture probabilities, could be another mechanism underlying the positive effect of lemmings on gulls hatching success. However, we recognize that the relationship between attack and capture probabilities and predator energetic balance was based on limited empirical data. Further refinement of the functions underlying changes in movement parameters and changes in attack and capture probabilities (Fig. 3 and Fig. 9) is needed to strengthen our conclusions of the relative importance of these mechanisms on gull hatching success. Overall, our study highlights how prey characteristics can lead to different mechanisms behind similar indirect interactions.

Although landscape characteristics are known to modulate the strength of predator-prey interactions (Barrios-O’Neill et al., 2015; Englund et al., 2011), few studies have explicitly included their effects in functional response models (see the review by Cherif et al. 2024). Our mechanistic model showed that nests located in different microhabitats generated variation in gull hatching success of 50% for the average lemming density. In the Arctic, wetland degradation is anticipated to accelerate with global warming, potentially leading to the formation of new refuges (islands), while some existing refuges may disappear (Corbeil-Robitaille et al., 2024). Although the net impact of these changes in refuge availability remains unknown, they could affect Arctic biodiversity by modulating the strength of predator-prey interactions. Thus, accurately anticipating ecosystem responses depends on reliable estimates of predation rates and on a mechanistic understanding of these interactions. As emphasized by Cherif et al. (2024) and DeLong (2021), the development of accurate models of predator-prey interactions is crucial for improving predictions of how food webs will respond to environmental changes.

## 6 Supplementary Methods

### 6.1 Parameter values used in the mechanistic model of predation

#### 6.1.1 Glaucous gull nest density

We calculated the average annual density of glaucous gull nests within within the home ranges of Arctic foxes. From 2008 to 2016, we quantified the annual home range of the fox from ARGOS telemetry data encompassing the gull monitoring area (*n* = 18 home ranges; data from Dulude et al. 2023). We estimated the average annual nest density at 0.26 nests per km^2^ (ranges from 0.05 to 0.54 nests per km^2^).

#### 6.1.2 Reaction distance

The reaction distance on a gull nest was defined as the distance at which the predator can initiate an attack. We used the same value as for the greater snow goose (*Anser caerulescens atlanticus*), since glaucous gulls are similar in mass and color. The body mass of the glaucous gull ranges from 1.25 kg to 2.7 kg and from 1.6 kg to 3.3 kg for the greater snow goose. This parameter was estimated from direct observations of foraging foxes on snow goose nests in summer 2019 (*µ* = 0.0328, se = 0.007, *n* = 25 attacks; Beardsell et al. 2021).

### 6.2 Parameter values of the energetic model

#### 6.2.1 Predator body mass, *M*

From 2004 to 2022, the body mass of adult foxes captured at our study site averaged 3.231kg (*sd* = 0.409, *n* = 420 foxes).

#### 6.2.2 Predator basal metabolic rate (BMR)

We used the equation presented in Careau et al. (2008) to estimate BMR based on the average fox mass (*M*):

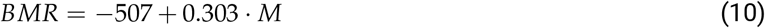

#### 6.2.3 Energetic content of prey

Brown lemming is the most abundant species at our study site, and adults weigh on average 51g (*sd* = 14 g, *n* = 1 450; from Seyer et al. 2020). The energetic content of a lemming was estimated at 6.276 KJ g^*−*1^ leading to 320 KJ for an average mass lemming. The energetic content estimate is based on the average of 220 animals (from 5 species of small mammals) analyzed with a calorimetric bomb (Górecki, 1965).

We estimated the energetic content of a sandpiper clutch (prey 2, *E*_2_) at 262 KJ per nest. The eggs of white-rumped and Baird’s sandpipers have a similar average mass of 9.32 g (Baird’s sandpiper: sd = 0.67, *n* = 30 eggs from 8 clutches (Moskoff and Montgomerie, 2020) and white-rumped sandpiper: *n* = 4 eggs from 1 clutch (Parmelee, 2020)). The energetic content was estimated at 7.03 KJ g^*−*1^ based on the energetic content of *Cortunix* quail eggs (a similar egg size; Prelipcean et al. 2012), which leads to 65.58 KJ per egg and 262 KJ per nest (clutch size of 4 eggs; Moskoff and Montgomerie 2020).

We estimated the energetic content of a glaucous gull clutch (prey 3, *E*_3_) at 1965.6 KJ (based on an average clutch size of 2.6 eggs). Glaucous gull eggs weigh on average 108 g (*n* = 31; Verboven et al. 2009) and their energetic content was estimated at 7 KJ g^*−*1^ based on the energetic content of herring gull (*Larus argentatus*) eggs (Hebert et al., 2020).

### 6.3 Sensitivity analysis

We conducted a local sensitivity analysis to evaluate how variations in attack and capture probabilities influence gull hatching success. Specifically, we simulated both linear and inverse sigmoidal relationships between attack and capture probabilities and the predator net energetic balance. We modified either the intercepts (for linear functions) or the asymptotes values (for sigmoidal functions) by *±*50% while keeping all others parameters constant, and assessed the resulting effects on gull hatching success. For instance, at a lemming density of 50 ind. per km^2^ (corresponding to a negative predator energetic balance), a *±*50% change in attack and capture probabilities intercepts (Fig. 7A1) leads to a variation in gull hatching success of 33% on shores and 29% on islands (Fig. 7A2). At a lemming density of 650 ind. per km^2^ (positive predator energetic balance), the same variation leads to a variation in gull hatching success of 74 % on shores and 21 % on islands (Fig. 7A2).

**Figure 7.**
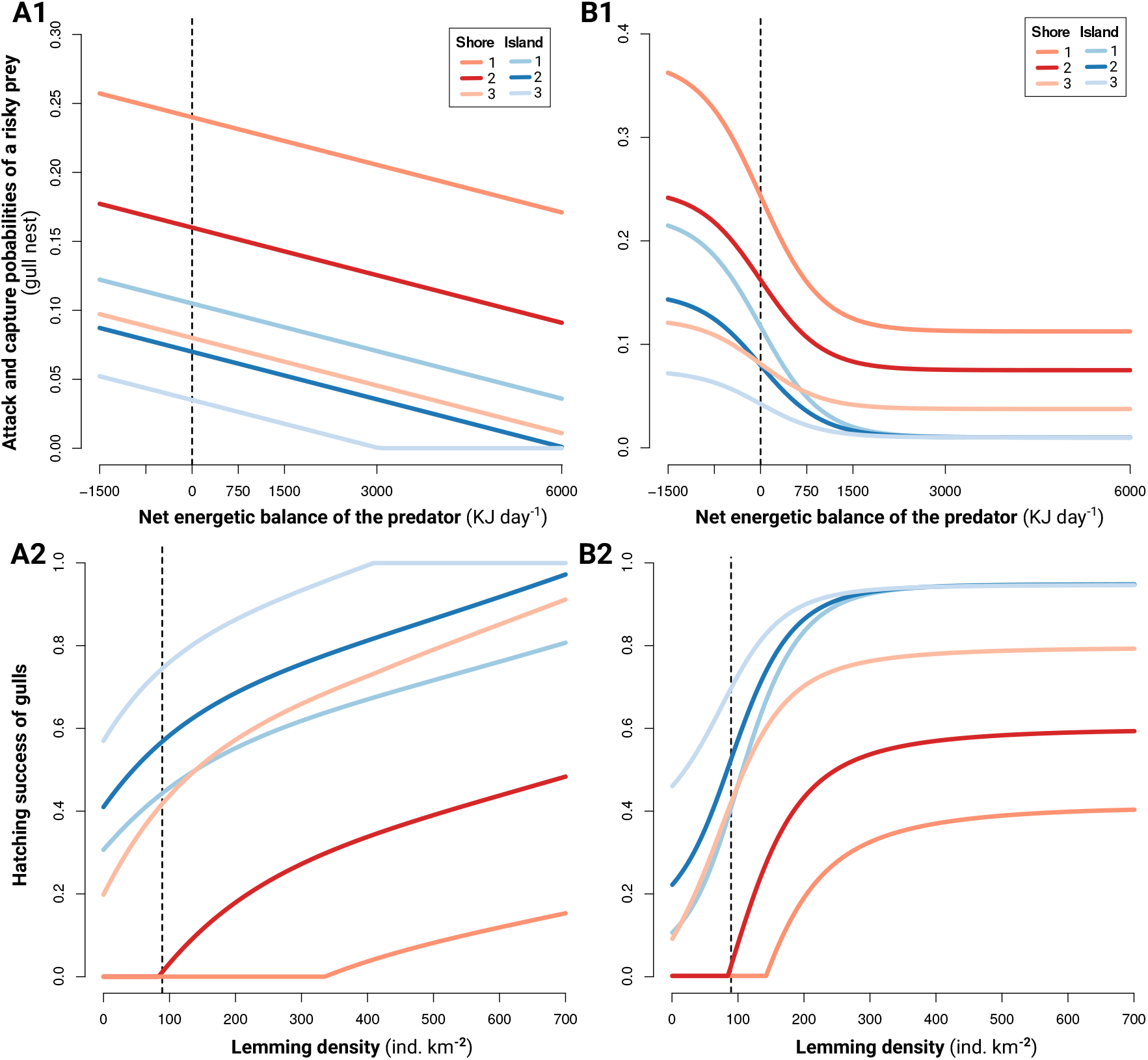
Simulations of linear (**A1**) and sigmoidal (**B1**) functions of attack and capture probabilities of a risky prey (gull nest) by the arctic fox in relation to the net energetic balance of the predator for the two habitats. Each colored line represents a different function. We modified the intercept (**A1**) and the asymptotes values (**B1**) of attack and capture probabilities by *±* 50% (relative to curve #2) while keeping the other parameters constant. (**A2-B2**) Relationship between hatching success of gull nests and lemming density in the two habitats predicted by the model based on the functions presented in **A1** and **B1**. Vertical black dashed line shows the threshold (predicted by the model) at which the net energetic balance of foxes changes from negative to positive.

**Figure 8.**
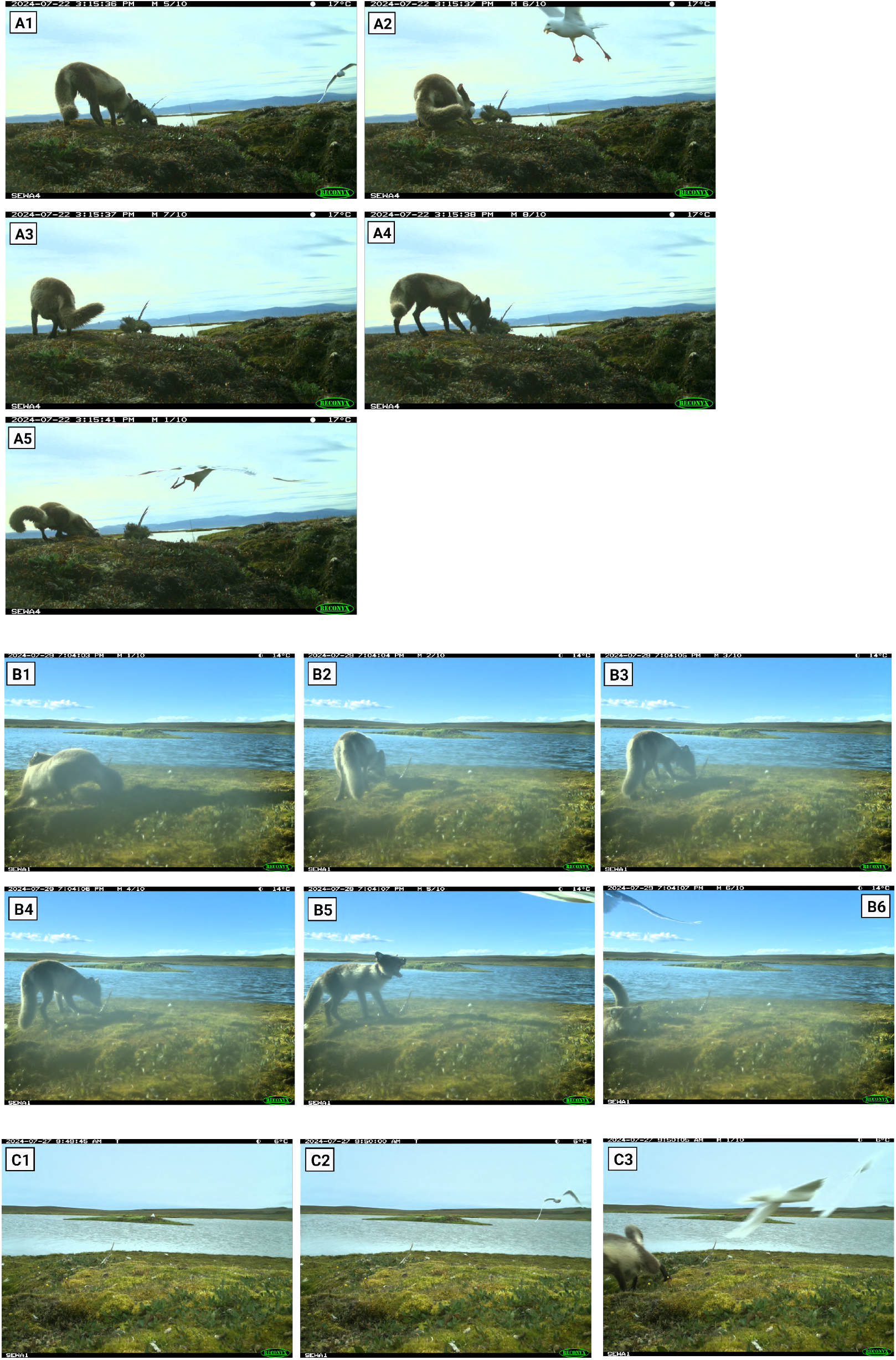
Image sequences (A, B, and C) of interactions between foxes and nesting glaucous gulls on Bylot Island in July 2024.

**Figure 9.**
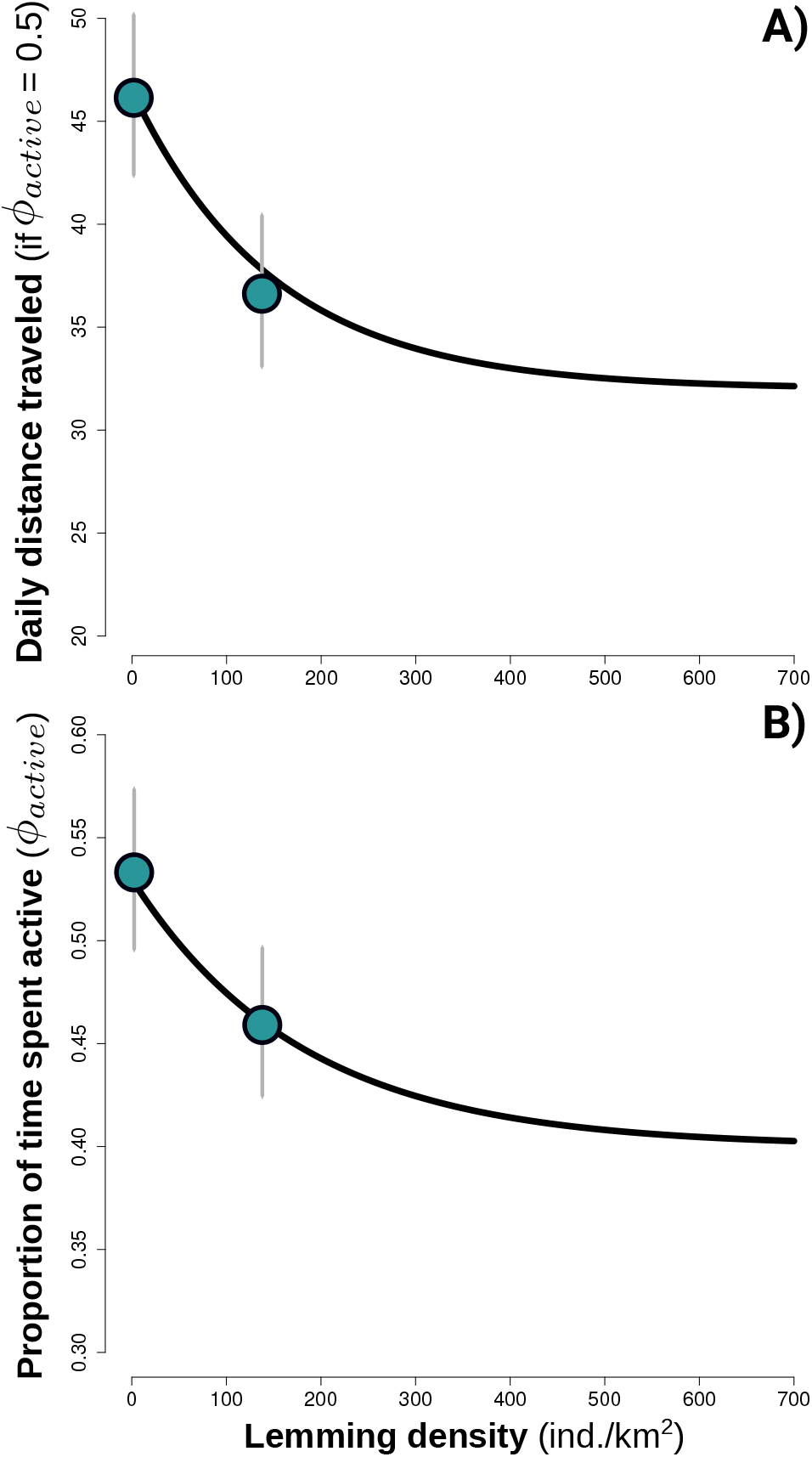
Relationships between lemming density and (A) the daily distance traveled by the arctic fox (when active), and (B) the proportion of time the Arctic fox spent active in a day. Both parameters are positively correlated. Points show the predicted proportion of time spent active per day and the predicted daily distance traveled by the fox between low and intermediate lemming densities (*n* = 371 fox-days). Error bars represent 95% confidence intervals. Relationships are from Beardsell et al. (2022) and are partially derived from empirical observations.

**Figure 10.**
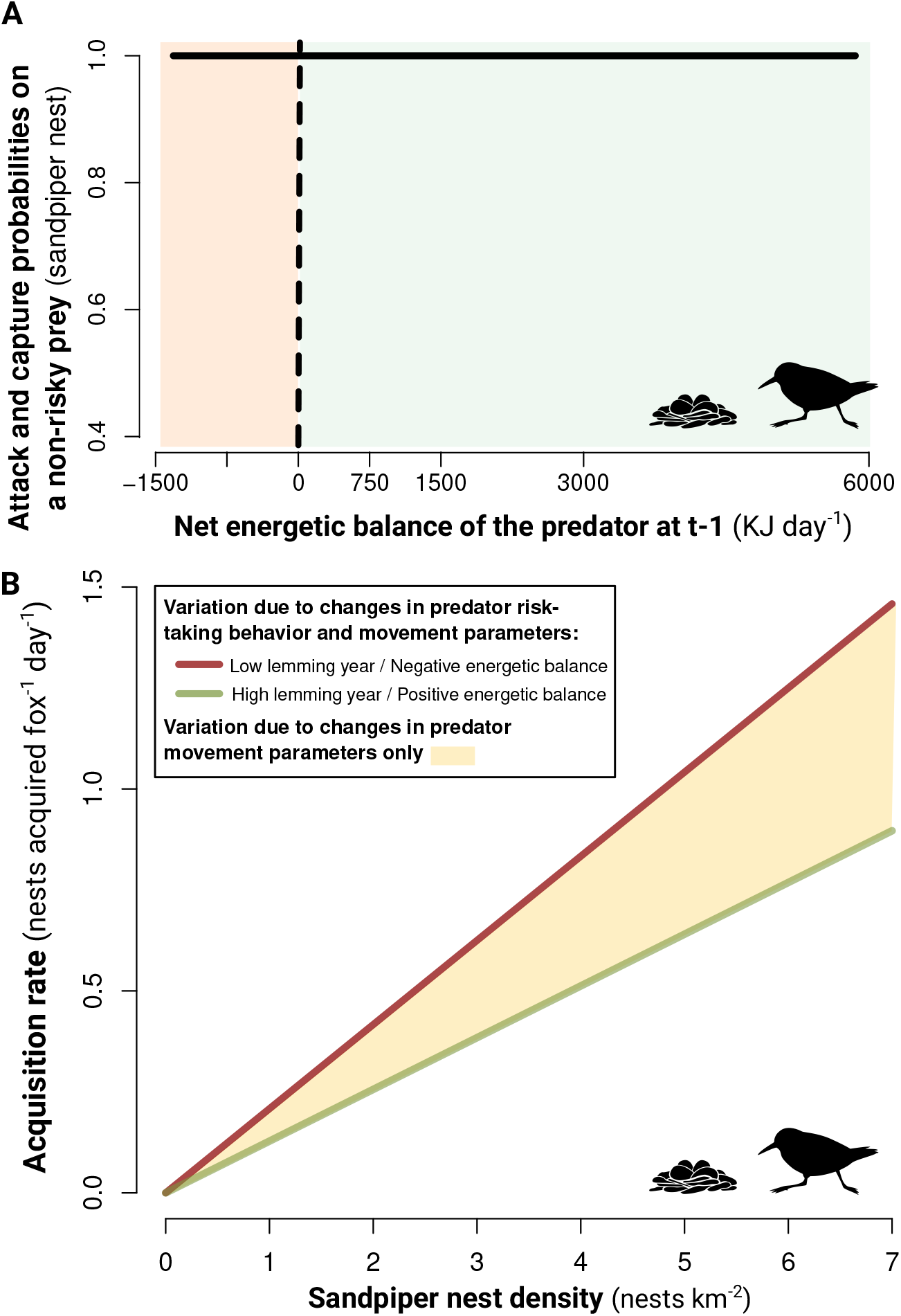
**A**) Relationship between daily attack and capture probabilities on a non-risky prey (sandpiper nests) by the Arctic fox at time t as a function of the net energetic balance of the predator at time t-1. Areas where the net energetic balance is >0 are green and <0 are red. **B)** Functional responses of the Arctic fox to sandpiper nests generated from the mechanistic multiprey model for a negative (red lines) and positive (green lines) energetic balance of the predator. Positive and negative energetic balance of the predator correspond, respectively, to lemming densities of 650 and 50 ind. per km^2^. The shaded area is based on a model without predator risk-taking behavior, retaining only changes in predator movement parameters (as a function of lemming density; Fig. S9). The density of gulls was set at the average value of 0.26 nests per km^2^.

## Acknowledgments

We are grateful to the many people who helped us with field work over the years, to the Mittimatalik Hunters and Trappers Organization, and to Park Canada staff for their assistance. The research relied on the logistic assistance of the Polar Continental Shelf Program (Natural Resources Canada) and of Sirmilik National Park of Canada. The research was funded by (alphabetical order): Arctic Goose Joint Venture, the ArcticNet Network of Centers of Excellence, the Canada Foundation for Innovation, the Canada Research Chairs Program, the Canadian Wildlife Service (Environment Canada), the Fonds de recherche du Québec-Nature et technologies, the International Polar Year program of Indian and Northern Affairs Canada, the Kenneth M Molson Foundation, the Natural Sciences and Engineering Research Council of Canada, the Northern Scientific Training Program, Polar Knowledge Canada, Université du Québec à Rimouski, and Université Laval.

## 7 Supplementary figures

## 8 Supplementary table

